# A new species of *Rana* from Anhui, China (Anura, Ranidae)

**DOI:** 10.64898/2026.04.24.720648

**Authors:** Zhirong He, Suyue Wang, Siyu Wu, Yufeng Bai, Jiedong Wei, Yingxin Li, Hongnuo Li, Yang Liu, Xiaohan Li, Xiaobing Wu, Supen Wang

## Abstract

The diversity of the brown frog genus *Rana* may be underestimated as the high similarity of morphological characters. A new species belonging to the genus *Rana* is delineated based on eight specimens obtained from the Tianma National Nature Reserve, Jinzhai County, Lu’an City, Anhui Province, China. The phylogenetic analysis based on three mitochondrial genes (12S, ND2, and Cyt b) and one nuclear gene (BDNF) showed that the new species formed an independent clade closely related to *R. culainensis* and received strong support. In addition, morphological differentiation confirmed the phylogenetic results, and both supported the validity of a new species (*Rana tianmaensis* sp. nov.) in the *R. japonica* species group. The discovery of this new species enhances people’s understanding of the biodiversity of *Rana* and can provide important foundational data for scientific decision–making on protected area construction, ecological conservation, and species diversity. With the inclusion of newly described species in this study, the distribution of *Rana* genus in China now includes 31 recognized species.

## 1. Introduction

The genus *Rana sensu lato* Linnaeus, 1758 ranks as the third most species–rich group within the family Ranidae, following Amolops Cope, 1865 (73 species) and Odorrana Fei, Ye & Huang, 1990 (62 species)[1]. Certain taxonomic perspectives propose that *Rana sensu lato* contains 114 species and 8 subgenera (*Rana, Amerana, Aquarana, Lithobates, Liuhurana, Pantherana, Pseudorana, Zweifelia*), thereby establishing it as the most extensive genus within the Ranidae family [2, 3]. In stark contrast, Frost (2025) [1] advocates for the elevation of the eight subgenera of *Rana* to full genus status, which would result in *Rana* (*sensu stricto*) comprising only 52 species. *Rana*, in its broad sense, has undergone two distinct dispersal events from East Asia to Northeastern and Northwestern North America, as well as a third dispersal event into Europe and Central Asia [4]. Recent scholarly inquiries into this genus have unearthed several novel species originating from China [5–10], indicating that the diversity of the genus is probably underestimated.

In the genus *Rana sensu lato*, thirty species have been documented within the Chinese region [2]. Through a meticulous examination of morphological characteristics, geographical distribution, and phylogenetic analysis, the subgenus *Rana* has been delineated into four species groups [4–6, 11–15]. These groups encompass (1) *R. japonica* group, twelve species: *R. chaochiaoensis* Liu, 1946, *R. chevronata* Hu & Ye, 1978, *R. culaiensis* Li, Lu & Li, 2008, *R. dabieshanensis, R. hanluica* Shen, Jiang & Yang, 2007, *R. japonica* Boulenger, 1879, *R. jiulingensis, R. jiemuxiensis* Yan, Jiang, Chen, Fang, Jin, Li, Wang, Murphy, Che & Zhang, 2011, *R. omeimontis* Ye & Fei, 1993, *R. longicrus* Stejneger, 1898, and *R. zhenhaiensis* Ye, Fei & Matsui, 1995, *R. zhijinensis* Luo, Xiao & Zhou, 2022; (2) *R. chensinensis* group, five species: *R. chensinensis* David, 1875, *R. dybowskii* Günther, 1876, *R. huanrenensis* Fei, Ye & Huang, 1990, *R. kukunoris* Nikolskii, 1918, and *R. taihangensis*; (3) *R. amurensis* group, three species: *R. amurensis* Boulenger, 1886, *R. coreana* Okada, 1928, and *R. luanchuanensis*; (4) *R. johnsi* group, three species: *R. johnsi* Smith, 1921, *R. sangzhiensis* Shen, 1986, and *R. wuyiensis*. Nonetheless, proposals for species groups have yet to be formulated to accommodate the remaining six species, i.e., *R. arvalis* Nilsson, 1842, *R. asiatica* Bedriaga, 1898, *R. maoershanensis* Lu, Li & Jiang, 2007, *R. sauter* Boulenger, 1909, *R. shuchinae* Liu, 1950, *R. minuscula* Liu, Wang, Huang, Liu, Yuan, and Che, 2024, while *R. weiningensis* [16] was replaced in the genus *Pseudorana* and recorded as *Pseudorana weiningensis* [16].

During our herpetological surveys in Jinzhai County, Lu’an City, Anhui Province, China, we collected a series of *Rana sensu lato* specimens. However, morphological examination showed that these specimens can be distinguished from its congeners. Subsequent molecular analyses corroborated this distinction that these specimens form an independent lineage within *Rana*. Consequently, in light of the disparities observed in both molecular and morphological characteristics, we suggest the population distributed in Jinzhai County, Lu’an City, Anhui Province, China as a new species, *Rana tianmaensis* sp. nov.

## 2. Materials and Methods

### 2.1. Sampling

Sampling procedures involving live *Rana* were in accordance with the Wild Animals Protection Law of China and approved by the Animal Ethics Committee at Anhui Normal University. A total of 16 specimens were collected from Anhui Province in this study: eight specimens of the describing species; eight of *R. zhenhaiensis*, from Huangshan City and Chizhou City (Figure 1). The specimens were humanely euthanized by immersion in a 95% ethanol solution, then transferred to 75% ethanol for preservation and storage. All these specimens were deposited in Anhui Normal University (ANU), Wuhu, Anhui.

**Figure 1.**
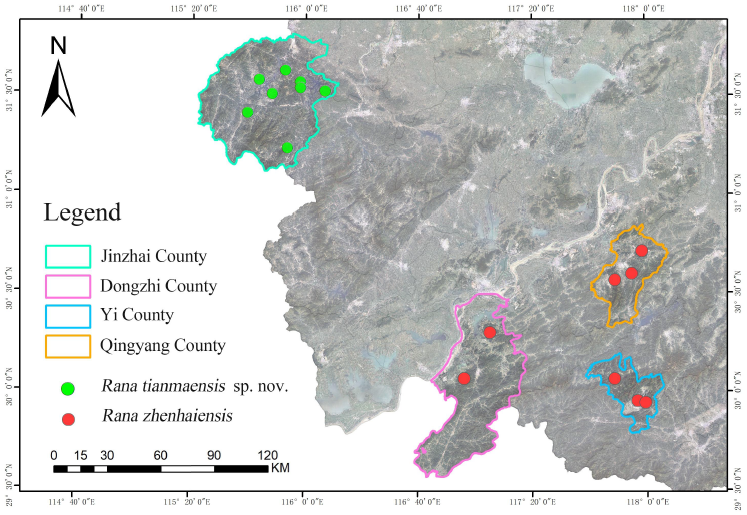
Sampling localities of *Rana tianmaensis* sp. nov. and *R. zhenhaiensis*.

### 2.2. DNA Extraction, PCR amplification, and sequencing

Genomic DNA was extracted from frozen or ethanol–preserved tissues using standard proteinase K extraction procedures [17]. Three mitochondrial genes and one nuclear genes were sequenced on sixteen samples we collected. The mitochondrial genes included 12S ribosomal RNA gene, NADH dehydrogenase subunit 2 (ND2) and cytochrome b (Cyt b). The nuclear DNA markers was brain–derived neurotrophic factor gene (BDNF). All primer sequences used for PCR and sequencing are summarized in Table 1. PCRs were performed with the same conditions in 25 µ L reaction volume: 20 to 80ng of genomic DNA, 25 µ L 2 × EasyTaq PCR SuperMix polymerase (TransGen Biotech, containing 1.25U Ex Taq, 0.4mM dNTP, 4mM Mg^2+^) and 0.4 µ M of primers. PCR amplifications were performed with the following cycling conditions: an initial denaturing step at 95 ° C for five min; 35 cycles of denaturing at 95 ° C for 30 s, 30 s at appropriate annealing temperature (Table 1) and extension at 72 ° C for 1 min; and a final extension step at 72 ° C for 10 min. The PCR products were purified using a rotating column and then sequenced using forward and reverse primers on the ABI Prism 3730XL automated DNA sequencer by Anhui General Biology Co., Ltd. All obtained sequences have been stored in GenBank (Table 2).

**Table 1.**
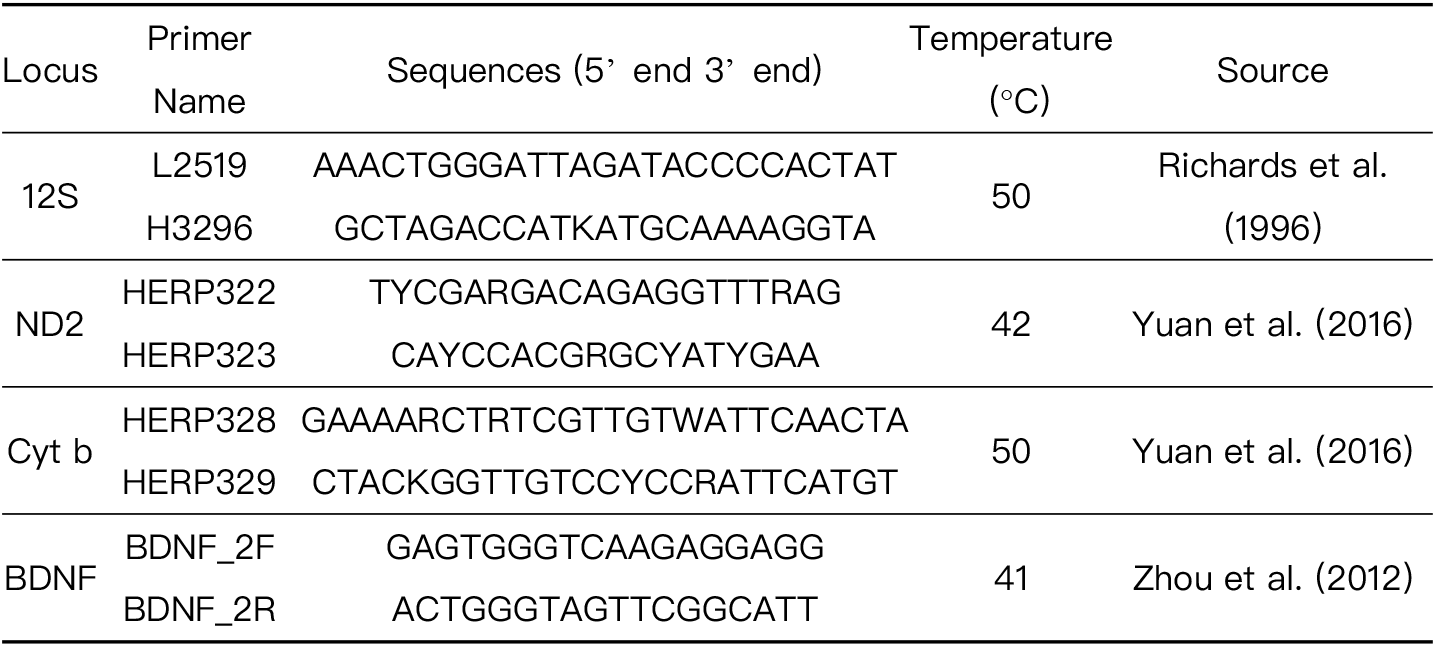
Primers used for PCR and sequencing.

**Table 2.**
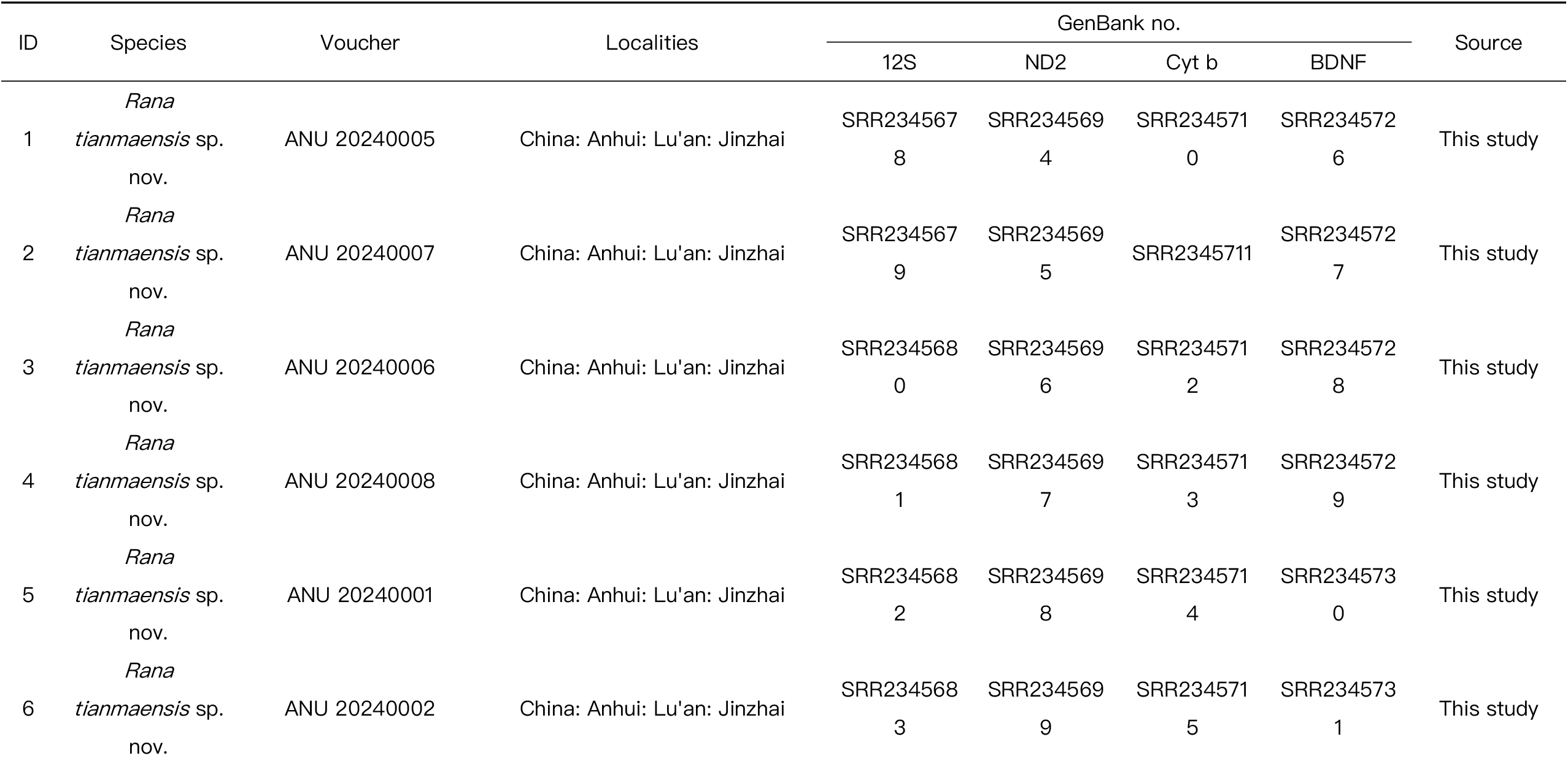
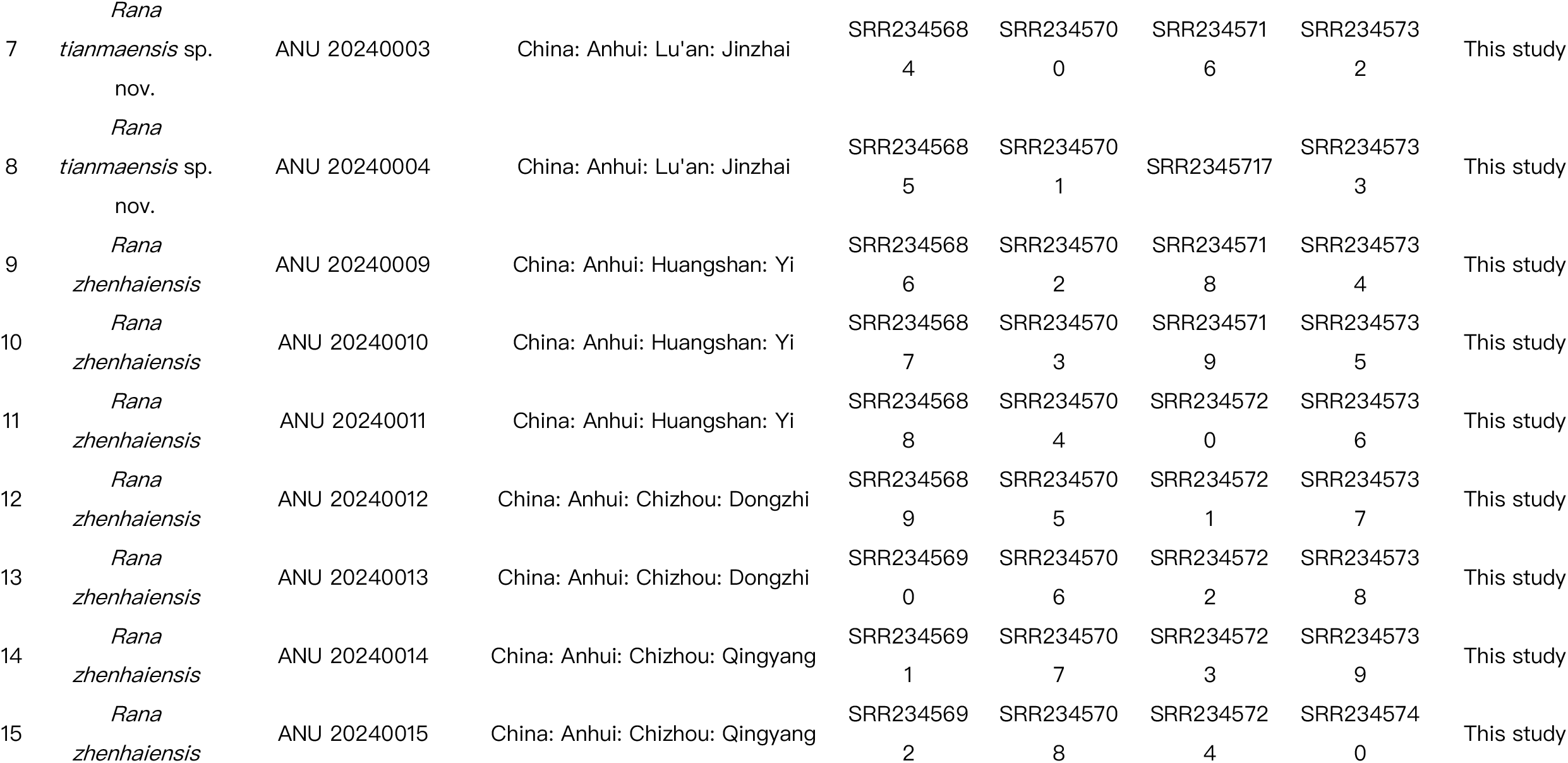
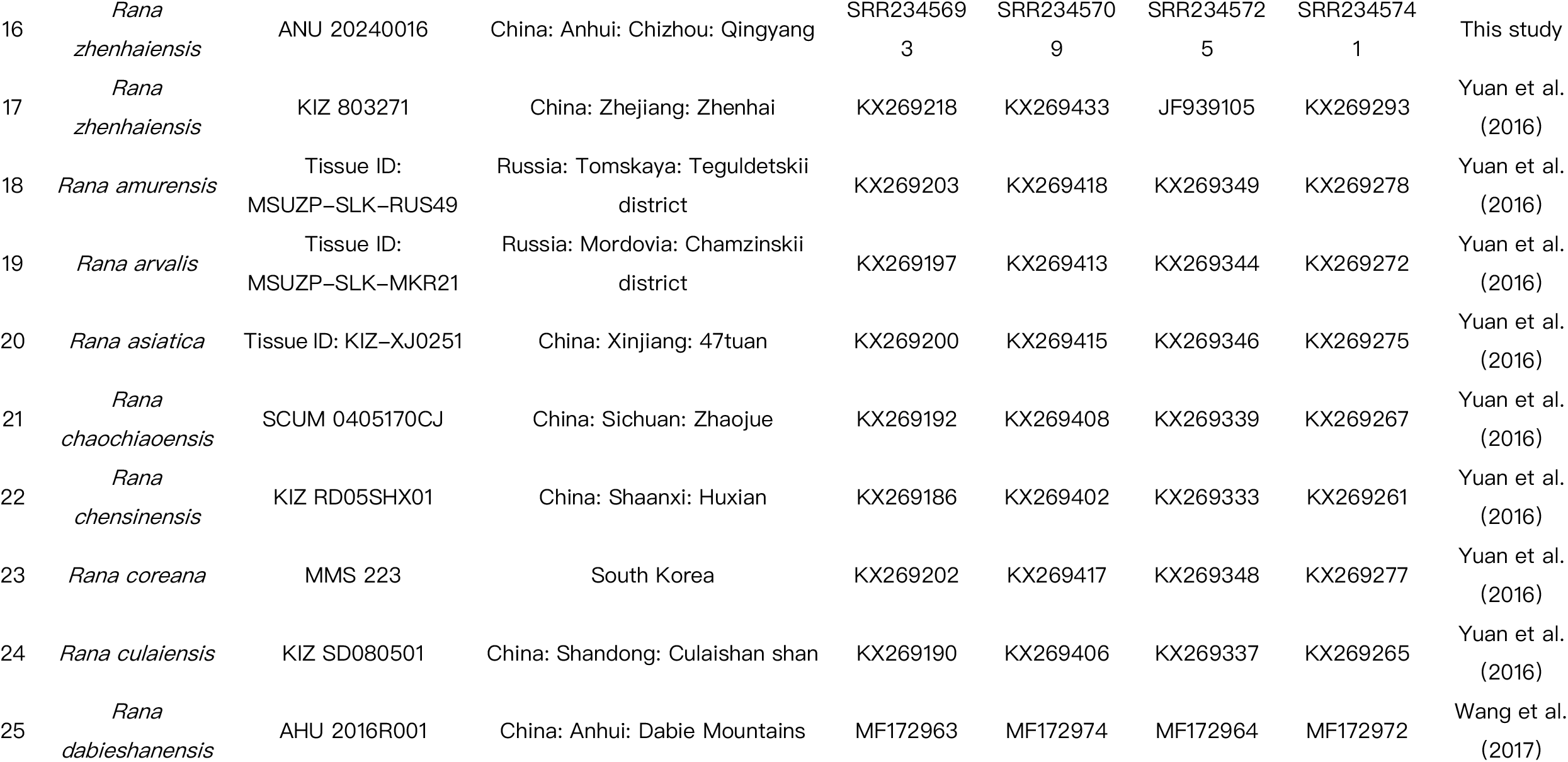
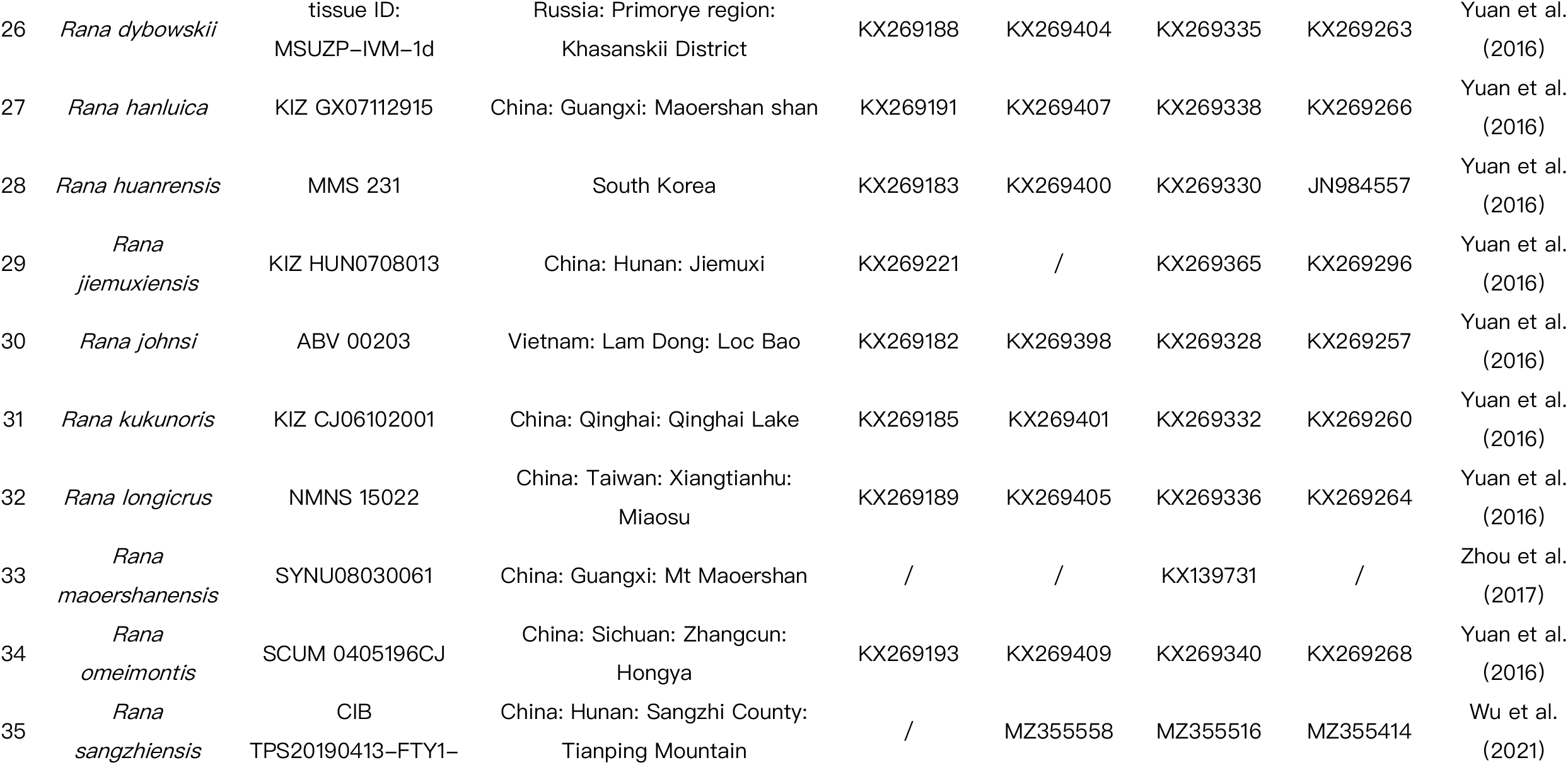
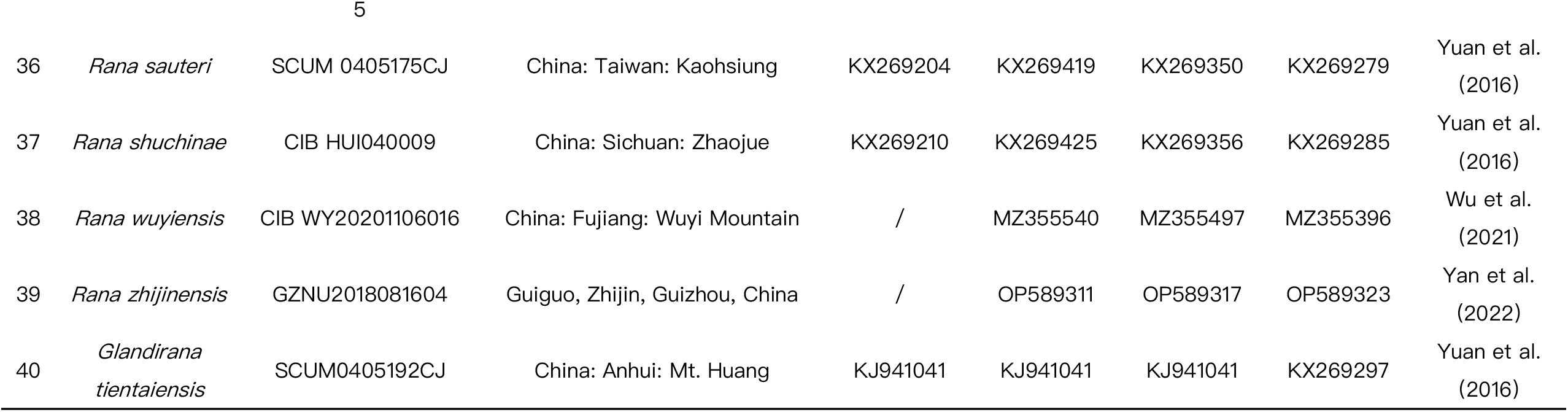
ID, Species, voucher museum numbers, sample localities and GenBank accession numbers for DNA sequences of *Rana* species used in the phylogenetic analyses.

### 2.3. Molecular phylogenetic analysis

In total, we retrieved 40 sequences from GenBank and included them in our phylogenetic analysis dataset, including 23 species of the genus *Rana* and one sequences of outgroups species from the genera *Glandirana* Fei, Ye, and Huang, 1990 (Table 2). The DNA sequences were aligned through the application of the Clustal W algorithm with default parameters [21]. The most suitable model for nucleotide evolution for each gene’s codon position and tRNA was ascertained via jModelTest 2.1.7 [22], based on the Akaike Information Criterion. The HKY+G+I model emerged as the optimal choice for each partition. The sequenced data were subjected to maximum likelihood (ML) analysis in MAGA12. In the ML analysis, a bootstrap consensus tree, derived from 1000 replicates was utilized to depict the evolutionary lineage of the analyzed taxa. We regarded tree topologies with bootstrap values (BS) of 70% or greater as sufficiently supported [23].

### 2.4. Morphological comparison and analyses

In morphological comparative analysis, the selected target traits can be broadly categorized two types: quantitative traits and qualitative traits. Quantitative traits refer to the continuous distribution of variation in a population of a certain trait, which rendering them indistinctly groupable. Quality traits refer to traits that can be qualitatively identified and clearly grouped within a population, with significant differences between groups.

#### Quantitative analysis

For the examination of quantitative traits, eight unnamed specimens alongside eight specimens of *R. zhenhaiensis* were externally measured utilizing a precision digital caliper (Deli DL91200 stainless steel 200mm Digita vernier caliper) with an accuracy of 0.01 millimeters. The morphological characteristics and their corresponding abbreviations, as detailed in [24] Table 3, while webbing formula adhered to [25]. Only adult specimens were measured. The determination of sex was based on secondary sexual characteristics.

**Table 3.** Abbreviations and descriptions of morphological characteristics of adult individuals..

| N<br>o. | Morphological<br>character | Abbrevi<br>ation | Description |
| --- | --- | --- | --- |
| 1 | Snout–vent<br>length | SVL | Distance from the tip of the snout to the articulation of<br>jaw |
| 2 | Head length | HL | Distance from the tip of the snout to the corner of the<br>mouth |
| 3 | Head width | HW | Maximum distance between the sides of the head |
| 4 | Snout length | SL | Distance from the tip of the snout to the front corner of<br>the eye |
| 5 | Internarial space | INS | Maximum distance between the inner edges of the left and right external nares |
| 6 | Diameter of eye | ED | Distance between the anterior and posterior corners of the eye |
| 7 | Interorbital space | IOS | Narrowest distance between the inner edges of the left and right upper eyelids |
| 8 | Upper eyelid width | UEW | Maximum width of the upper eyelid |
| 9 | Tympanum diameter | TD | Maximum diameter of the tympanum |
| 10 | Lower arm and hand length | LAHL | Distance from the elbow to the distal end of the Finger III |
| 11 | Hand length | HAL | Distance from the base of the hand to the tip of Toe III |
| 12 | Diameter of lower arm | LAD | Maximum diameter of the lower arm |
| 13 | Hindlimb length | HLL | Distance from the middle part of the posterior end of the body to the tip of the Toe IV |
| 14 | Tibia length | TL | Distance between the two ends of the tibia |
| 15 | Tibia width | TW | The greatest width of tibia |
| 16 | Foot length | FL | Distance from distal end of shank to the tip of Toe IV |
| 17 | Length of foot and tarsus | TFL | Distance from the tibiotarsal articulation to the distal end of the Toe IV |

#### Qualitative analysis

We employed the aforementioned seventeen morphological features to assign ratings to each individual. During comparisons, only those traits ubiquitous among all population members were taken into account, while traits exhibiting variation among individuals were excluded, in order to delineate discrete and diagnostic attributes (following Wiens 2000 criteria).

## 3. Results

### 3.1. Molecular phylogenetic relationships

This study conducted a meticulous phylogenetic analysis of mitochondrial and nuclear gene fragments derived from eight unnamed specimens, eight *R. zhenhaiensis*, 23 recognized species belonging to the genus *Rana*, and one known outgroup species *Glandirana tientaiensis*. The nucleotide sequences, employed to reconstruct the phylogenetic tree subsequent to alignment, encompassed mitochondrial 12S (448 bp), ND2 (706 bp), Cyt b (513 bp), and the nuclear gene BDNF (367 bp).

The phylogenetic tree reconstructed utilizing both mitochondrial and nuclear gene datasets exhibited topological discrepancies when compared to the tree reconstructed solely using mitochondrial gene datasets (Figure 2)., but all support *R. johnsi* group as a basal branch in the four species groups, *R. johnsi, R. chensinensis, R. amurensis*, and *R. japonica*, within the subgenus *Rana*. Both phylogenetic trees indicate that *R. japonica* group is the sister group of *R. chensinensis* group. The P–distances based on the mitochondrial and nuclear gene datasets among species are presented in Table 4 and Table 5. Within the *R. johnsi* group, two major branches are discernible. One of the branches is composed of *R. chaochiaoensis* and *R. zhijinensis*. Another branch shows two branches. One of the branches is composed of *R. jiemuxiensis*. Another branch showed two branches, but the bootstrap support values for both phylogenetic trees were not high (BS < 70). One of the branches is composed of *R. hanluica*. Another branch showed two branches, but the bootstrap support values for both phylogenetic trees were not high (BS < 70). One of the branches is composed of *R. omeimontis* and *R. dabieshanensis* (BS 92). Another branch shows three branches (BS 99). The first branch is composed of *R. zhenhaiensis*. The second branch is composed of *R. longicirus*. The third branch consists of *R. culaiensis* and the unnamed samples from Jinzhai County, Lu’an City, Anhui Province, China. The unnamed samples collected in Jinzhai formed a unique lineage (BS 94). The results of both maximum likelihood (ML) analyses were largely congruent, as illustrated in Figure 2, with the *Rana* samples from Jinzhai County, Lu’an City forming a highly supported sister species lineage with *R. culaiensis* (BS 98).

**Table 4.** Mean p–distance (%) of three mitochondrial genes (12S, ND2 and Cyt b) and one nuclear gene (BDNF) among the genus *Rana* species used in this study.

| ID | Species | 1 | 2 | 3 | 4 | 5 | 6 | 7 | 8 | 9 | 10 | 11 | 12 | 13 | 14 | 15 | 16 | 17 | 18 | 19 | 20 | 21 | 22 | 23 | 24 |
| --- | --- | --- | --- | --- | --- | --- | --- | --- | --- | --- | --- | --- | --- | --- | --- | --- | --- | --- | --- | --- | --- | --- | --- | --- | --- |
| 1 | *Rana tianmaensis* sp. nov. | — |  |  |  |  |  |  |  |  |  |  |  |  |  |  |  |  |  |  |  |  |  |  |  |
| 2 | *Rana zhenhaiensis* | **0.0379** |  |  |  |  |  |  |  |  |  |  |  |  |  |  |  |  |  |  |  |  |  |  |  |
| 3 | *Rana amurensis* | **0.2706** | 0.2524 |  |  |  |  |  |  |  |  |  |  |  |  |  |  |  |  |  |  |  |  |  |  |
| 4 | *Rana arvalis* | **0.2605** | 0.2429 | 0.1171 |  |  |  |  |  |  |  |  |  |  |  |  |  |  |  |  |  |  |  |  |  |
| 5 | *Rana asiatica* | **0.2591** | 0.2424 | 0.1126 | 0.0928 |  |  |  |  |  |  |  |  |  |  |  |  |  |  |  |  |  |  |  |  |
| 6 | *Rana chaochiaoensis* | **0.2481** | 0.2251 | 0.1176 | 0.1181 | 0.1167 |  |  |  |  |  |  |  |  |  |  |  |  |  |  |  |  |  |  |  |
| 7 | *Rana chensinensis* | **0.2674** | 0.2460 | 0.1236 | 0.1137 | 0.1118 | 0.1084 |  |  |  |  |  |  |  |  |  |  |  |  |  |  |  |  |  |  |
| 8 | *Rana coreana* | **0.2614** | 0.2463 | 0.0914 | 0.1285 | 0.1186 | 0.1246 | 0.1276 |  |  |  |  |  |  |  |  |  |  |  |  |  |  |  |  |  |
| 9 | *Rana culaiensis* | **0.2061** | 0.1897 | 0.1230 | 0.1161 | 0.1122 | 0.0816 | 0.1207 | 0.1220 |  |  |  |  |  |  |  |  |  |  |  |  |  |  |  |  |
| 10 | *Rana dabieshanensis* | **0.2272** | 0.2078 | 0.1201 | 0.1151 | 0.1082 | 0.0875 | 0.1222 | 0.1285 | 0.0514 |  |  |  |  |  |  |  |  |  |  |  |  |  |  |  |
| 11 | *Rana dybowskii* | **0.2633** | 0.2430 | 0.1241 | 0.1142 | 0.1117 | 0.1177 | 0.0989 | 0.1305 | 0.1196 | 0.1241 |  |  |  |  |  |  |  |  |  |  |  |  |  |  |
| 12 | *Rana hanluica* | **0.2287** | 0.2083 | 0.1156 | 0.1181 | 0.1067 | 0.0811 | 0.1137 | 0.1260 | 0.0558 | 0.0558 | 0.1196 |  |  |  |  |  |  |  |  |  |  |  |  |  |
| 13 | *Rana huanrensis* | **0.2733** | 0.2505 | 0.1300 | 0.1206 | 0.1122 | 0.1172 | 0.0440 | 0.1290 | 0.1275 | 0.1251 | 0.0900 | 0.1196 |  |  |  |  |  |  |  |  |  |  |  |  |
| 14 | *Rana jiemuxiensis* | **0.0446** | 0.0446 | 0.0848 | 0.0886 | 0.0749 | 0.0606 | 0.0804 | 0.0787 | 0.0409 | 0.0409 | 0.0788 | 0.0409 | 0.0848 |  |  |  |  |  |  |  |  |  |  |  |
| 15 | *Rana johnsi* | **0.2585** | 0.2434 | 0.1250 | 0.1161 | 0.1117 | 0.1295 | 0.1261 | 0.1280 | 0.1201 | 0.1210 | 0.1246 | 0.1255 | 0.1369 | 0.0772 |  |  |  |  |  |  |  |  |  |  |
| 16 | *Rana kukunoris* | **0.2703** | 0.2494 | 0.1246 | 0.1127 | 0.1028 | 0.1058 | 0.0376 | 0.1241 | 0.1196 | 0.1216 | 0.0865 | 0.1127 | 0.0331 | 0.0841 | 0.1280 |  |  |  |  |  |  |  |  |  |
| 17 | *Rana longicrus* | **0.2172** | 0.1946 | 0.1245 | 0.1225 | 0.1136 | 0.0835 | 0.1246 | 0.1265 | 0.0193 | 0.0573 | 0.1211 | 0.0598 | 0.1290 | 0.0477 | 0.1220 | 0.1216 |  |  |  |  |  |  |  |  |
| 18 | *Rana maoershanensis* | **0.1572** | 0.1662 | 0.1621 | 0.1523 | 0.1309 | 0.1543 | 0.1680 | 0.1797 | 0.1602 | 0.1484 | 0.1621 | 0.1563 | 0.1777 | 0.1621 | 0.1680 | 0.1582 | 0.1719 |  |  |  |  |  |  |  |
| 19 | *Rana omeimontis* | **0.2348** | 0.2141 | 0.1161 | 0.1201 | 0.1082 | 0.0924 | 0.1241 | 0.1304 | 0.0553 | 0.0296 | 0.1246 | 0.0568 | 0.1280 | 0.0447 | 0.1220 | 0.1246 | 0.0563 | 0.1738 |  |  |  |  |  |  |
| 20 | *Rana sangzhiensis* | **0.3176** | 0.2956 | 0.1524 | 0.1354 | 0.1316 | 0.1519 | 0.1468 | 0.1505 | 0.1410 | 0.1467 | 0.1486 | 0.1493 | 0.1588 | 0.0968 | 0.0323 | 0.1486 | 0.1423 | 0.1738 | 0.1461 |  |  |  |  |  |
| 21 | *Rana sauteri* | **0.2594** | 0.2409 | 0.1126 | 0.1181 | 0.0973 | 0.1251 | 0.1157 | 0.1201 | 0.1161 | 0.1206 | 0.1201 | 0.1117 | 0.1186 | 0.0757 | 0.1285 | 0.1092 | 0.1196 | 0.1465 | 0.1225 | 0.1581 |  |  |  |  |
| 22 | *Rana shuchinae* | **0.2677** | 0.2520 | 0.1270 | 0.1230 | 0.1165 | 0.1359 | 0.1340 | 0.1299 | 0.1354 | 0.1339 | 0.1256 | 0.1314 | 0.1280 | 0.0901 | 0.1245 | 0.1300 | 0.1408 | 0.1699 | 0.1369 | 0.1455 | 0.1285 |  |  |  |
| 23 | *Rana wuyiensis* | **0.3124** | 0.2928 | 0.1474 | 0.1328 | 0.1208 | 0.1513 | 0.1513 | 0.1480 | 0.1347 | 0.1385 | 0.1455 | 0.1423 | 0.1619 | 0.0923 | 0.0342 | 0.1505 | 0.1385 | 0.1719 | 0.1373 | 0.0310 | 0.1461 | 0.1385 |  |  |
| 24 | *Rana zhijinensis* | **0.3021** | 0.2764 | 0.1392 | 0.1328 | 0.1271 | 0.0297 | 0.1304 | 0.1442 | 0.0911 | 0.0999 | 0.1309 | 0.0968 | 0.1347 | 0.0706 | 0.1550 | 0.1265 | 0.0980 | 0.1680 | 0.1069 | 0.1575 | 0.1366 | 0.1493 | 0.1543 | — |

**Table 5.**
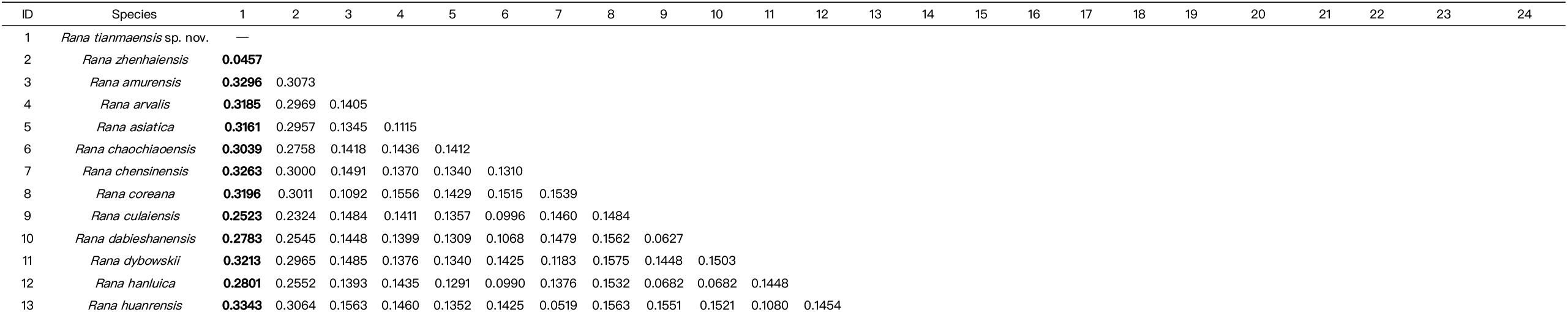
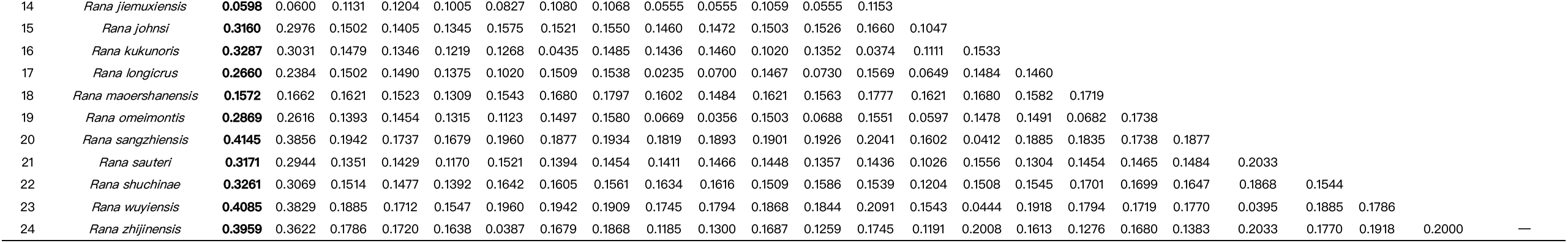
Mean p–distance (%) of three mitochondrial genes (12S, ND2 and Cyt b) among the genus *Rana* species used in this study.

**Figure 2.**
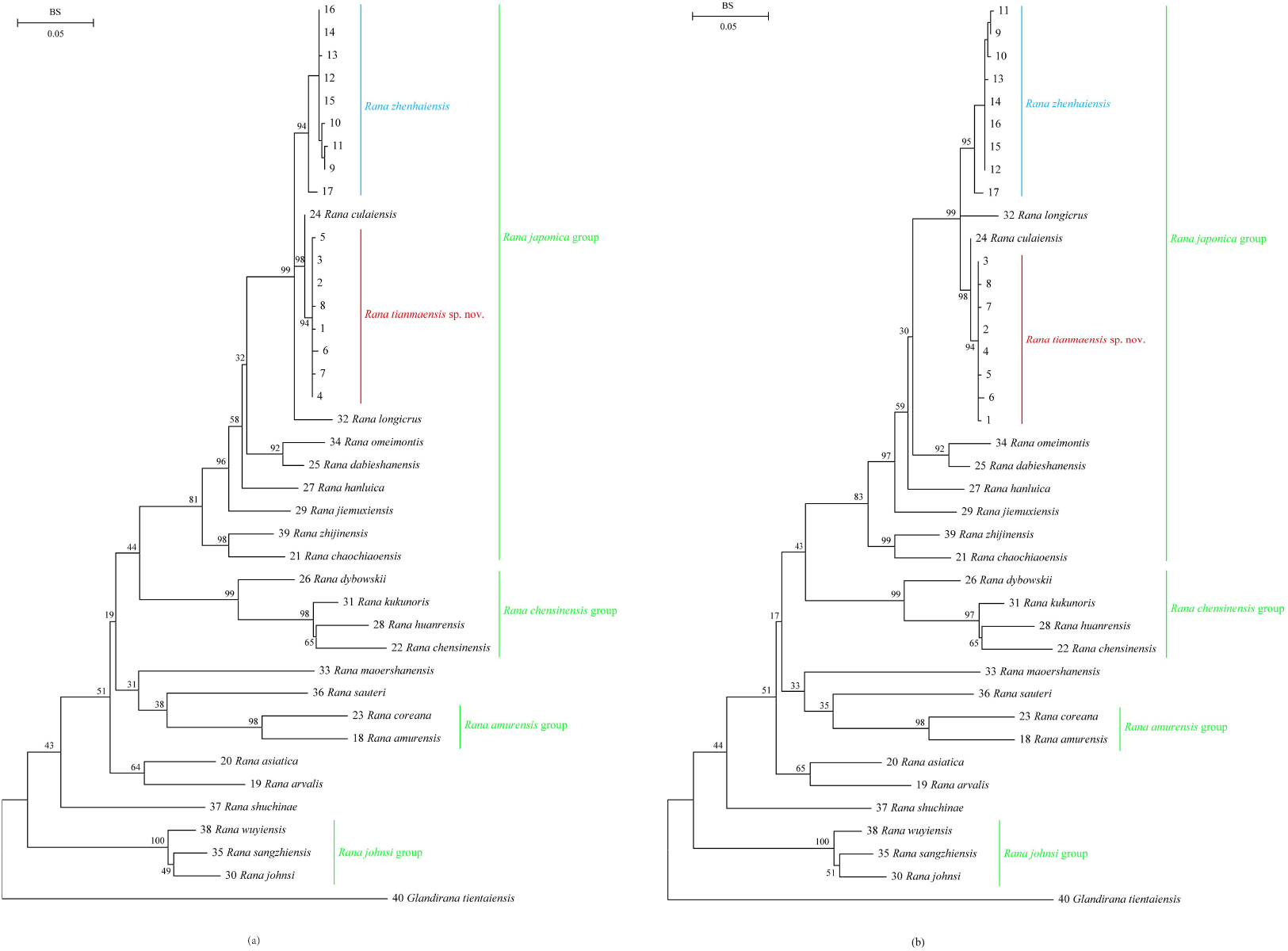
Phylogenetic relationships of *Rana tianmaensis* sp. nov. and its relatives. (a) Maximum likelihood (ML) tree constructed based on three mitochondrial genes (12S, ND2 and Cyt b) and one nuclear gene (BDNF); (b) Maximum likelihood (ML) tree constructed based on three mitochondrial genes (12S, ND2 and Cyt b). In both phylogenetic tree, bootstrap supports (BS) values from ML analyses are given beside nodes. Scale bars denote nucleotide substitutions per sites for mitochondrial and nuclear genes.

### 3.2. Morphological comparisons

Morphologically, these eight specimens of the *Rana* from unnamed lineages can be distinguished from all known recognized reliably (with detailed taxonomic accounts provided below). The unnamed *Rana* population from Jinzhai County, Lu’an City and the sister taxon *R. zhenhaiensis* from Yi County, Dongzhi County, and Qingyang County were statistically analyzed by morphometric measurements (Table 6; Figure 3). The results of ANOVA on morphometrics for males revealed significant differences between individuals of the unnamed *Rana* population and *R. zhenhaiensis* in SVL, HL, SL, IOS, LAHL, HAL, HLL, TL, FL and TFL for males (*P*–values < 0.05). The results of ANOVA on morphometrics for females showed that individuals of the unnamed *Rana* population and *R. zhenhaiensis* are significantly different in SVL, HL, HW, SL, IOS, UEW, LAHL, HAL, LAD, HLL, TL, TW, FL and TFL for females (*P*–values < 0.05). For the PCA result of males, the first two principal components (PCs) account for 99.4% and 0.3% of the variance, respectively (in total > 90.0% of the variance). For the PCA result of females, the first two principal components (PCs) account for 95.2% and 3.4% of the variance, respectively (in total > 90.0% of the variance). As illustrated in the scatter plots (Figure 3), the individuals of the unnamed *Rana* population and *R. zhenhaiensis* formed respective clusters and were clearly separated.

**Table 6.**
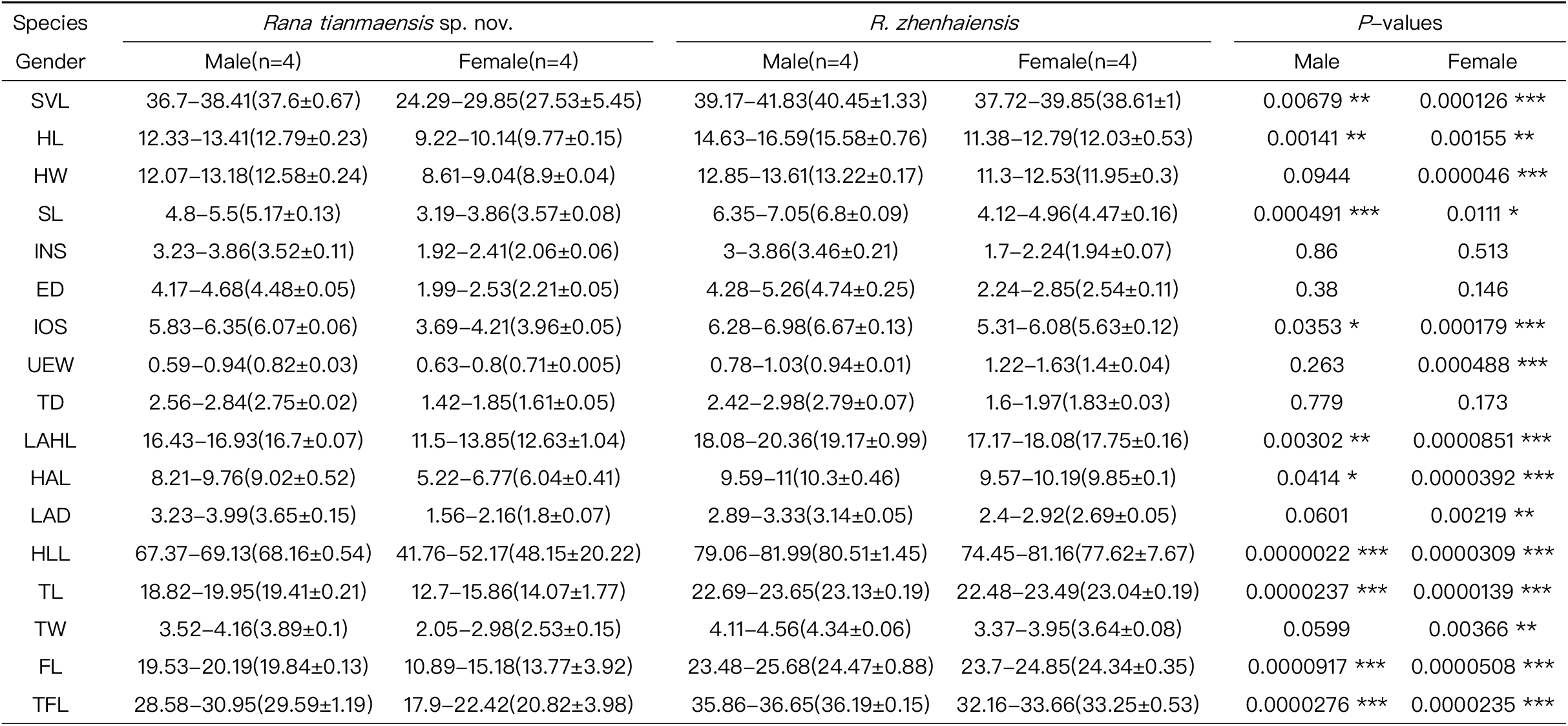
Measurements (in mm) of the adult specimens of *Rana tianmaensis* sp. nov. and *R. zhenhaiensis*. See abbreviations for the morphological characters in the materials and methods section.

**Figure 3.**
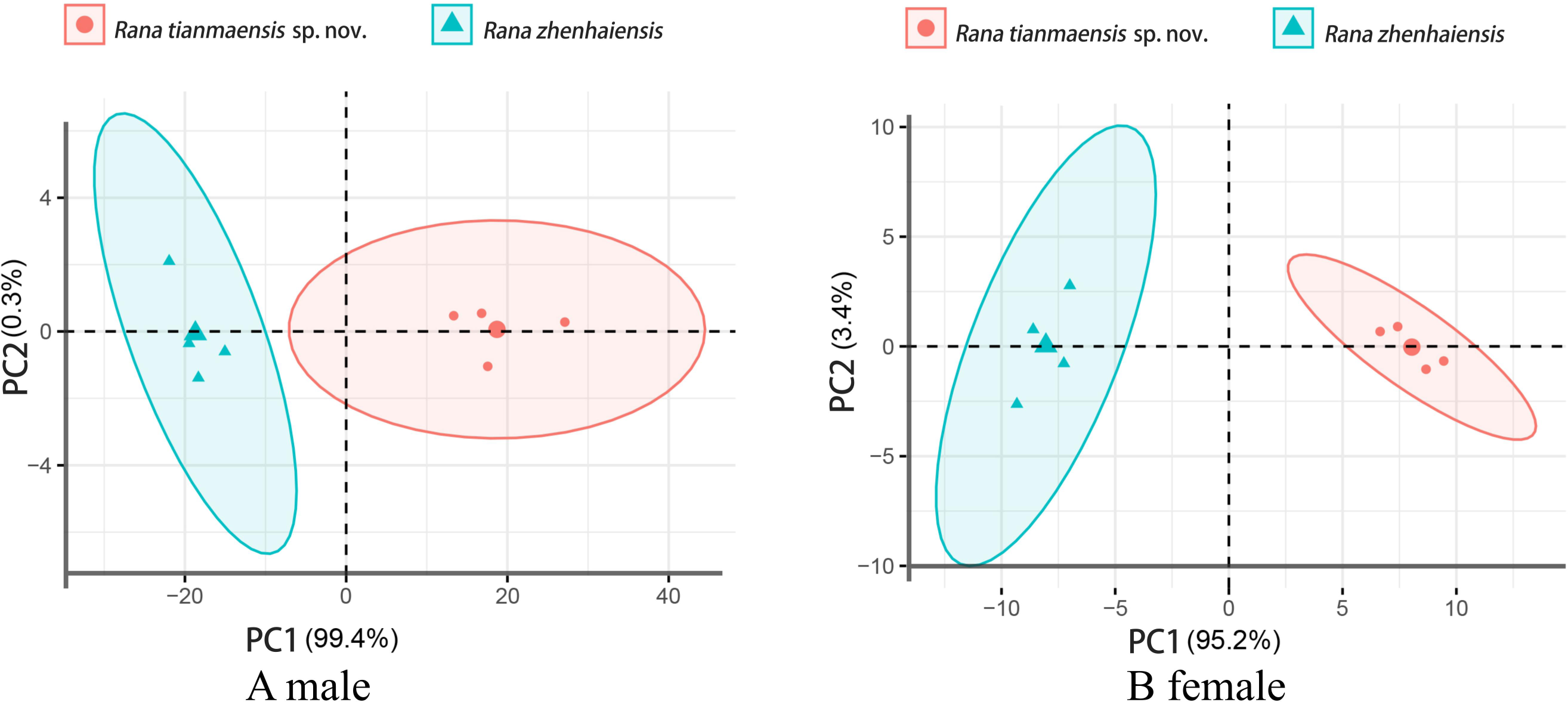
Scatter plot of PC1 and PC2 of Principal Component Analysis based on the morphometric measurements, distinguishing the samples of *Rana tianmaensis* sp. nov. and *R. zhenhaiensis*. The slightly larger point in the middle displays the midpoint of the group.

Therefore, combined with the results above morphological examination, morphometric statistical analysis and phylogenetic analysis, we describe the specimens collected from Jinzhai County as a new species, *Rana tianmaensis* sp. nov.

### 3.3. Taxonomic accounts

#### Holotype

ANU 20240001, an adult male, collected by Zhirong He and Yufeng Bai on 14 June 2024, in a small pond at an altitude of 672 meters in Tianma National Nature Reserve (31.29916282 ° N, 115.68566238 ° E), Jinzhai County, Lu’an City, Anhui Province, China.

#### Paratypes

Three adult males: ANU 20240002 – ANU 20240004, four adult females: ANU 20240005 – ANU 20240008, collected by Zhirong He and Yufeng Bai from 15 to 26 July 2024, in Jinzhai County, Lu’an City, Anhui Province, China.

#### Etymology

Tianma National Nature Reserve, located in Jinzhai County, Anhui Province, is centered around two provincial–level nature reserves, Tiantangzhai and Mazongling. It is a nature reserve of forest ecosystem type. Tianma National Nature Reserve is rich in biological resources and has strong representativeness and typicality. This valuable biological gene bank plays an important role in maintaining the ecological environment and promoting sustainable development of the ecological economy, with high protection value and comprehensive benefits. Meanwhile, all unnamed specimens we collected this time were sourced from within the Tianma National Nature Reserve. Therefore, we suggest naming the unnamed *Rana* species discovered this time after the Tianma National Nature Reserve, which is commonly known in Latin as “*Rana tianmaensis*” and in Chinese as “天马林蛙( tiān mǎ lín wā)”.

#### Diagnosis

(1) a small–sized frog of the genus *Rana*,, SVL = 36.7–38.41 mm in adult males (n = 4), 24.29–29.85 mm in adult females (n = 4); (2) head length slightly larger than head width, HL/HW 1.02 in males and 1.10 in females; (3) internarial space significantly smaller than interorbital space, INS/IOS 0.58 in males and 0.52 in females; (4) dorsolateral fold present and thin, extending straight from tympanum to above the groin; (5) tympanum diameter significantly smaller than the diameter of eye, TD/ED = 0.61 in males and 0.73 in females; (6) fingers circummarginal grooves and webbed absent, relative finger lengths I < II < IV < III; (7) toes circummarginal grooves absent, toe webbing present, relative toe lengths I < II < V < III < IV; (8) breeding males possess dark gray–blackish nuptial pad separated into three parts on finger I; (9) tibiotarsal articulation reaching far forward beyond tip of the snout.

#### Description of the holotype

ANU 20240001 (Figure 4), adult male, SVL 38.16 mm; head slightly compressed vertically, length slightly longer than width (HL/HW 1.11); snout projecting forward, somewhat pointed at tip; nostril lateral, closer to tip of snout than eye; canthus rostralis distinct; eye moderate (ED/HL ratio 0.33); internasal space less than interorbital space (INS/IOS ratio 0.65); tympanum distinct, vertically oval, slightly longer than half of eye diameter (TD/ED ratio 0.61), tympanic rim not elevated; pupil horizontally oval.

**Figure 4.**
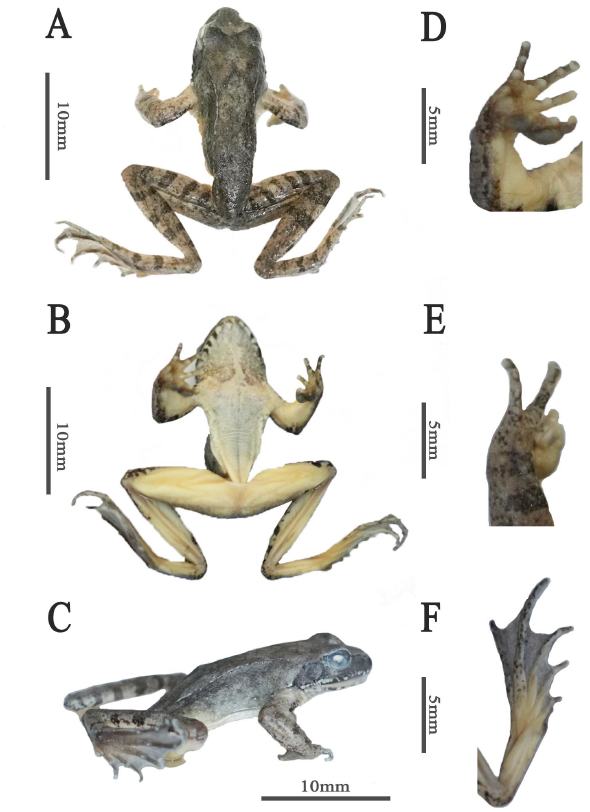
Morphological features of the adult male holotype ANU 20240001 of *Rana tianmaensis* sp. nov. **A** dorsal view. **B** ventral view **C** lateral view **D** ventral view of hand **E** dorsal view of hand **F** ventral view of foot.

Forelimb robust; length of lower arm and hand, less than body length (LAHL/SVL ratio 0.44); fingers slender, without webbing; tips of fingers rounded but not expended; relative finger length I<II<IV<III; subarticular tubercles distinct; three metacarpal tubercles, inner one largest and rounded, outer one smallest and oval, middle one long, oval; inner metacarpal tubercle indistinct, ovoid, mostly covered by nuptial pad; nuptial pad on finger I, divided into three parts, prior two parts completely joined together.

Hindlimb moderately long, tibiotarsal articulation reaching anterior corner of eye, heels overlap when flexed and held perpendicular to body; tibia length slightly larger than half of body length (TL/SVL ratio 0.51). Tips of all toes rounded and slightly compressed, not expanded; relative toe length I<II<V<III<IV; webbing on toes developed, nearly entire web; inner metatarsal tubercle elongate, oval; outer metatarsal tubercle rounded, smaller than the inner one.

Skin relatively smooth; dorsal surface of head smooth, dorsal surface of body scattered with a few tubercles; lateral surface of body with small tubercles, slightly less than dorsal body; dorsolateral fold narrow and slightly curved, extending from temporal fold above tympanum fold to groin, not connecting to posterior corner of eye; longitudinal skin ridges on dorsal tibia distinct; throat, chest, and belly of thighs being smooth.

#### Coloration in life

(Figure 5) Dorsal parts of head and dorsum, flank, forelimb, thigh, tibia, and foot brownish, with four and three black transverse bands on dorsal thigh and tibia, respectively; loreal region brownish; black spots below canthus rostralis; maxillary gland brownish–black; tympanic region yellowish–brown; lateral side of body brown, discontinuous and thin black line below dorsolateral fold; ventral surface of head yellowish–white, with dense and small brownish–black spots; ventral body yellowish–white, small brownish–black spots scattered on throat; an distinct inverted V–shaped black glandular ridge in scapular region and extending backward to middle of dorsal surface of body to form a distinct inverted V–shaped black glandular ridge at posterior end; nuptial pad dark gray–blackish.

**Figure 5.**
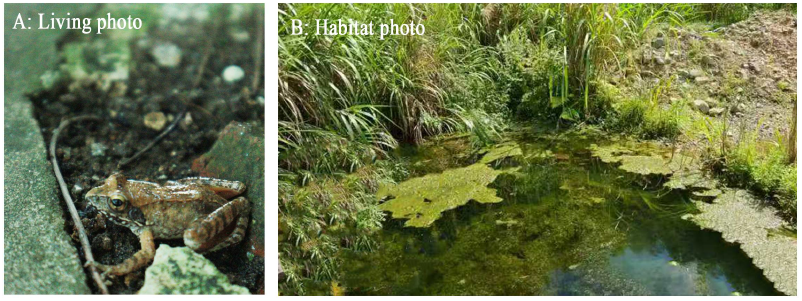
A: *Rana tianmaensis* sp. nov. in life. B: Habitat at the *Rana tianmaensis* sp. nov. in Tianma National Nature Reserve, Jinzhai, Anhui, China.

#### Sexual dimorphism

Forearm of males much stronger than females (LAD male/female=2.02); well–developed nuptial pad present in males, pale gray in coloration.

#### Distribution

Currently known only from the type locality.

#### Variation

All measurements of adult specimens are given in Table 6.

#### Comparisons

The new species is assigned to the *R. japonica* group based on the following morphological characteristics, i.e., digits without circummarginal grooves, and dorsolateral fold distinct, extending straight from temporal fold above tympanum to above the groin. Here, the morphology of the new species with 11 species of the *R. japonica* group were compared.

The new species is phylogenetically closest to *R. culaiensis*, but can be morphologically distinguished from the latter by combining the following morphological characteristics: (1) the head length of *Rana tianmaensis* sp. nov. is slightly larger than its head width, while the head length of *R. culaiensis* is slightly less than its head width; (2) *Rana tianmaensis* sp. nov. has an inverted V–shaped mark on its back surface, while *R. culaiensis* does not; (3) the breeding males of *Rana tianmaensis* sp. nov. have dark gray black nuptial pad, while the breeding males of *R. culaiensis* have dark brown nuptial pad; (4) the tibia of male *Rana tianmaensis* sp. nov. is slightly shorter than their feet, while the tibia of male *R. culaiensis* is slightly longer than their feet.

Compared to the eleven remaining species in the *R. japonica* group, the results are as follows.

*Rana tianmaensis* sp. nov. and *R. zhenhaiensis* can be distinguished by the following morphological features (Table 6): (1) the relative finger length of *Rana tianmaensis* sp. nov. is I<II<IV<III, while the relative finger length of *R. zhenhaiensis* is II<IV<I<III; (2) the relative finger length of *Rana tianmaensis* sp. nov. is I < II < V < III < IV, while the relative finger length of *R. zhenhaiensis* is I < II < V = III < IV.

*Rana tianmaensis* sp. nov. and *R. chaochiaoensis* can be distinguished by the following morphological features [11]: (1) the head length of *Rana tianmaensis* sp. nov. is slightly larger than its head width, while the head length of *R. chaochiaoensis* is smaller than its head width; (2) the relative finger length of *Rana tianmaensis* sp. nov. is I<II<IV<III, while the relative finger length of *R. chaochiaoensis* is II<IV<I<III; (3) the relative toe length of *Rana tianmaensis* sp. nov. is I < II < V < III < IV, while the relative toe length of *R. chaochiaoensis* is I < II < V = III < IV; (4) The nuptial pad of the breeding male of *Rana tianmaensis* sp. nov. is dark gray black, divided into three parts, while that of *R. chaochiaoensis* is gray, divided into four parts.

*Rana tianmaensis* sp. nov. and *R. chevronta* can be distinguished by the following morphological features [10]: (1) the internarial distance of *Rana tianmaensis* sp. nov. is significantly smaller than interorbital distance, while the internarial distance of *R. chevronta* is larger than the interorbital distance; (2) the relative finger length of *Rana tianmaensis* sp. nov. is I<II<IV<III, while the relative finger length of *R. chevronta* is II<IV<I<III; (3) the nuptial pad of the breeding male of *Rana tianmaensis* sp. nov. is dark gray black, divided into three parts, while that of *R. chevronta* is purplish gray, undivided, and complete.

*Rana tianmaensis* sp. nov. and *R. dabieshanensis* can be distinguished by the following morphological features [14]: (1) the relative finger length of *Rana tianmaensis* sp. nov. is I<II<IV<III, while the relative finger length of *R. dabieshanensis* is II<IV<I<III; (2) *Rana tianmaensis* sp. nov. has an inverted V-shaped mark on its back surface, while *R. dabieshanensis* does not; (3) the nuptial pad of the breeding male of *Rana tianmaensis* sp. nov. divided into three parts, while that of *R. dabieshanensis* divided into two parts.

*Rana tianmaensis* sp. nov. and *R. hanluica* can be distinguished by the following morphological features [26]: (1) the relative finger length of *Rana tianmaensis* sp. nov. is I<II<IV<III, while the relative finger length of *R. hanluica* is II < I < IV < III; (2) the nuptial pad of the breeding male of *Rana tianmaensis* sp. nov. is dark gray black, divided into three parts, while that of *R. hanluica* is gray, divided into two parts.

*Rana tianmaensis* sp. nov. and *R. japonica* can be distinguished by the following morphological features [10]: the nuptial pad of the breeding male of *Rana tianmaensis* sp. nov. is dark gray black, divided into three parts, while that of *R. japonica* is grayish brown or yellowish brown and divided into two parts.

*Rana tianmaensis* sp. nov. and *R. jiulingensis* can be distinguished by the following morphological features [5]: (1) the head length of *Rana tianmaensis* sp. nov. is slightly larger than its head width, while the head length of *R. jiulingensis* is significantly smaller than its head width; (2) the internarial distance of *Rana tianmaensis* sp. nov. is significantly smaller than interorbital distance, while the internarial distance of *R. jiulingensis* is significantly larger than the interorbital distance; (3) *Rana tianmaensis* sp. nov. has an inverted V-shaped mark on its back surface, while *R. jiulingensis* does not; (4) the nuptial pad of the breeding male of *Rana tianmaensis* sp. nov. is dark gray black, while that of *R. jiulingensis* is creamy white.

*Rana tianmaensis* sp. nov. and *R. jiemuxiensis* can be distinguished by the following morphological features [13]: (1) the head length of *Rana tianmaensis* sp. nov. is slightly larger than its head width, while the head length of *R. jiemuxiensis* is slightly smaller than its head width; (2) *Rana tianmaensis* sp. nov. has an inverted V-shaped mark on its back surface, while *R. jiemuxiensis* does not; (3) the relative finger length of *Rana tianmaensis* sp. nov. is I<II<IV<III, while the relative finger length of *R. jiemuxiensis* is II<IV<I<III; (4) the nuptial pad of the breeding male of *Rana tianmaensis* sp. nov. is dark gray black, divided into three parts, while that of *R. jiemuxiensis* is gray, divided into two parts.

*Rana tianmaensis* sp. nov. and *R. omeimontis* can be distinguished by the following morphological features [27]: (1) *R. omeimontis* has a larger body size than *Rana tianmaensis* sp. nov.; (2) *Rana tianmaensis* sp. nov. has an inverted V-shaped mark on its back surface, while *R. omeimontis* does not; (3) the head length of *Rana tianmaensis* sp. nov. is slightly larger than its head width, while the head length of *R. omeimontis* is slightly smaller than its head width; (4) the relative finger length of *Rana tianmaensis* sp. nov. is I<II<IV<III, while the relative finger length of *R. omeimontis* is II<IV<I<III; (5) the relative toe length of *Rana tianmaensis* sp. nov. is I < II < V < III < IV, while the relative toe length of *R. omeimontis* is I < II < V = III < IV.

*Rana tianmaensis* sp. nov. and *R. longicrus* can be distinguished by the following morphological features [10]: the relative finger length of *Rana tianmaensis* sp. nov. is I<II<IV<III, while the relative finger length of *R. longicrus* is II < I < IV < III.

*Rana tianmaensis* sp. nov. and *R. zhijinensis* can be distinguished by the following morphological features [8]: (1) the body size of *Rana tianmaensis* sp. nov. is smaller than that of *R. zhijinensis*; (2) the head length of *Rana tianmaensis* sp. nov. is slightly larger than its head width, while the head length of *R. zhijinensis* is equal to its head width; (3) the internarial distance of *Rana tianmaensis* sp. nov. is significantly smaller than interorbital distance, while the internarial distance of *R. zhijinensis* is equal to the interorbital distance; (4) the dorsal lateral folds of *Rana tianmaensis* sp. nov. extend straight from tympanum to above the groin, while the dorsal lateral folds of *R. zhijinensis* straight from posterior margin of the upper eyelid to above the groin; (5) the relative toe length of *Rana tianmaensis* sp. nov. is I < II < V < III < IV, while the relative toe length of *R. zhijinensis* is I < II < III < V < IV.

*Rana tianmaensis* sp. nov. and *R. chensinensis* group can be distinguished by the following morphological features [7, 11]: the nuptial pad of the breeding male of *Rana tianmaensis* sp. nov. is dark gray black, divided into three parts, while that of *R. chensinensis* group divided into four parts, except in *R. taihangensis* (divided into two parts).

## 4. Discussion

Because of the phenotypic plasticity and local morphological adaptations to the similar habitats, cryptic species are prevalent among amphibians [35–37]. Recently, the integrative taxonomy of morphological comparisons and molecular analyses has greatly facilitated the identification of numerous cryptic amphibian species [36, 38–41].

The Tianma National Nature Reserve in Anhui Province is situated in Jinzhai County, Lu’an City, at the junction of Hubei, Henan, and Anhui provinces. The continuous mountain ranges within its boundaries, coupled with a favorable natural environment, make it an ideal habitat for *Rana*. Eight specimens (ANU20240001 – ANU20240008) sampled from Tianma National Nature Reserve were also subjected to detailed morphological and molecular studies pertaining to Chinese amphibians.

The phylogenetic analysis results reveal that the specimens discovered in the Tianma National Nature Reserve have evolved into a unique lineage (BS 94) and exhibit a close affinity with *R. culaiensis*. The formation of independent branches between different species is unequivocally established. So, the specimens from Tianma National Nature Reserve ought to be classified as a novel species within the genus *Rana*. The reconstructed phylogenetic tree exhibits topologies that diverge from those reported in recent studies [7, 8, 10], but all support the monophyly of *R. japonica* group, *R. chensinensis* group, *R. amurensis* group, and *R. johnsi* group within the subgenus *Rana* (Figure 2). This inconsistent topology may stem from variations in the number and type of genetic markers employed, as well as the evolutionary models used, as different genes may have different rates of evolution [28]and phylogenetic analyses may yield conflicting results. Therefore, phylogenetic studies using a certain number of mitochondrial and nuclear genes can significantly contribute to elucidating the phylogeny and taxonomy of the genus *Rana* [4]. In addition, morphological differentiation corroborates the phylogenetic findings, and collectively reinforcing the validity of the newly proposed species (*Rana tianmaensis* sp. nov.) in the *R. japonica* species group.

Numerous phylogeographic studies have demonstrated that mountain ranges can serve as geographic barriers and thereby facilitating speciation [29–32]. At the species level, “subdivision” is advisable. This approach aids in a more effective and precise understanding and description of the natural history of species, as well as fostering coherent communication and actions in taxonomy and conservation biology practices [33]. Given the phylogenetic and morphological disparities, it is reasonable to consider unnamed populations as independent species. With the inclusion of newly described species in this study, the distribution of *Rana* genus in China now includes 31 recognized species.

Currently, three species of the genus *Rana* sensu lato are distributed in Jinzhai County, Lu’an City, Anhui Province, namely *R. dabieshanensis, R. zhenhaiensis, R. chensinensis* [34]. However, the intricate geographical distribution of these species remains unclear. A definitive species diversity inventory can serve as a pivotal source of fundamental data for making scientific decisions pertaining to protected area establishment, ecological preservation, and species diversity conservation. Therefore, we advocate for increased efforts to protect the Tianma National Nature Reserve and ascertain the habitat of the newly discovered species *Rana tianmaensis* sp. nov. to facilitate further research on its systematic evolution and biogeography.

## 5. Conclusions

In summary, we have described a new species of *Rana* genus from Anhui, China, i.e., *Rana tianmaensis* sp. nov., based on molecular and morphological evidence. The new species formed an independent clade closely related to *R. culainensis* and supported the validity of a new species (*Rana tianmaensis* sp. nov.) in the *R. japonica* species group. With the inclusion of newly described species in this study, the distribution of *Rana* genus in China now includes 31 recognized species. It is crucial to increase sampling and obtain more molecular data in order to better understand the phylogenetic and taxonomic membership relationships of the *Rana* genus.

## Author Contributions

Miss Zhirong He: Conceptualization (Lead); Data curation (Lead); Software (Lead); Methodology (Lead); Writing – original draft (Lead). Miss Suyue Wang: Data curation (Equal); Supervision (Equal); Validation (Equal); Writing – review & editing (Equal). Miss Siyu Wu: Data curation (Equal); Software (Equal); Formal analysis (Equal); Supervision (Equal); Visualization (Equal). Mr. Yufeng BAI: Investigation (Equal); Supervision (Equal); Writing – review & editing (Equal). Mr. Jiedong WEI: Investigation (Equal); Supervision (Equal); Writing – review & editing (Equal). Miss Yingxin Li: Investigation (Equal); Supervision (Equal); Writing – review & editing (Equal). Miss Hongnuo Li: Investigation (Equal); Supervision (Equal); Writing – review & editing (Equal). Miss Yang Liu: Investigation (Equal); Supervision (Equal); Writing – review & editing (Equal). Miss Xiaohan Li: Investigation (Equal); Supervision (Equal); Writing – review & editing (Equal). Prof. Xiaobing WU: Funding acquisition (Equal); Resources (Equal); Supervision (Equal); Validation (Equal); Writing – review & editing (Equal). Prof. Supen WANG (Corresponding Author): Funding acquisition (Equal); Resources (Equal); Supervision (Equal); Validation (Equal); Writing – review & editing (Equal).

## Data Availability Statement

All the sequences used in this study were accessed through the GenBank database, and the accession numbers are listed in Table 2. Morphological specimens were deposited at Anhui Normal University (ANU), Wuhu, Anhui.

## Acknowledgments

We thank anonymous reviewers for their valuable suggestions on this work.

